# siRNA Mediated Genetic Perturbation of Primary Human Leukemia Stem and Progenitor Cells

**DOI:** 10.1101/2025.09.17.676969

**Authors:** Anagha Inguva, Hunter Tolison, Jeremy Rahkola, Craig T. Jordan, Courtney Jones, Maria Amaya

**Affiliations:** Division of Hematology, University of Colorado School of Medicine, Aurora, CO, USA; Rocky Mountain Regional VAMC, Aurora CO; Division of Experimental Hematology and Cancer Biology, Cincinnati Children’s Hospital Medical Center, Cincinnati, OH, USA

## Abstract

Acute myeloid leukemia (AML) and myelodysplastic syndromes (MDS) are aggressive hematologic malignancies with poor outcomes. Leukemia stem and progenitor cells (LSPCs) are a subset of cells within the bulk tumor thought to be responsible for initiating disease and causing relapse. LSPCs evade chemotherapy partially due to their quiescent state. Therefore, studying this cell subpopulation is critical to identify new disease targets that can better the outcomes of AML/MDS patients. The use of RNAi and genetic approaches is technically challenging in primary leukemia cells and particularly LSPCs. Overcoming this technical hurdle could greatly expand the breath of pre-clinical and mechanistic examination of LSPCs. In this study, we describe a methodology to efficiently introduce siRNA, resulting in effective gene knock down in LSPCs. After isolation of LSPCs from primary patient samples, we showed that electroporation of these cells does not affect cell viability significantly. Further, using siGLO green transfection indicator, we show that RNA is introduced into the nucleus of these cells. siRNA transfection leads to efficient knockdown of target genes and has subsequent biological relevant activity. We have identified a method to effectively knock down genes in leukemia stem and progenitor cells, opening up new avenues to examine LSPC biology in human specimens.

## Introduction

Acute myeloid leukemia (AML) and myelodysplastic syndromes (MDS) are aggressive malignancies with poor outcomes.^1,2^ These diseases are characterized by abnormal differentiation and clonal expansion of myeloid progenitors. A subpopulation of these malignant cells called leukemia stem cells (LSCs) are a critical component of disease progression and their abundance correlates with disease aggressiveness.^3,4^ LSCs are thought to be more quiescent, and therefore more difficult to eliminate with standard therapy. Therefore, studying ways to target these cells is critical in the field of myeloid malignancies.

Our laboratory and others have shown various methods to isolate leukemia stem cells in order to perform detailed characterization of LSCs.^5-7^ However, genetically manipulating these cells can be challenging given they replicate less frequently than bulk leukemia cells and they have limited viability in culture. In this study, we have built upon an electroporation method to genetically modify primary human and mouse hematopoietic cells,^8^ and we have optimized a method of obtaining efficient knockdown in LSPCs, which leads to significant biological effects.

## Methods

### Transfection/Electroporation

For transfections, we used the Neon Transfection System (Invitrogen). Cells spun and re-suspended in buffer T and 50nM non-targeting siRNA (Dharmacon) at 2 million cells per 100uL of Buffer T according to the manufacturer’s protocol. Electroporation settings were as follows: 1,600 V, 10ms, 3 pulses. Cells were then plated in IMDM-based media.

### Flow Cytometry/Image Stream

Viability was measured using Dapi and measured by flow cytometry. siGLO green activity was measured by flow cytometry using GFP positivity within the DAPI-cell population. For ImageStream studies, FIX & PERM cell fixation assay (ThermoFisher) was used to stain for intracellular marker Dapi. Cells were then washed with FACS Buffer (1% FBS in 1X PBS) and stained with the extracellular maker CD34 for MDS cells. Cells were then run on the ImageStreamX MkII (Cytek Biosciences). Samples were then analyzed using the Ideas 6.2 software.

### Engraftment Assays

Leukemia stem cell function was assessed by measuring engraftment of primary AML specimens post transfection with either siRNA scramble or siMYC in one experiment and siGLO+ and siGLO-in the other experiment. NSG-S mice were conditioned with 25 mg/kg busulfan via i.p. injection. Second day at injection, primary AML cells post transfection were cultured for 1-2 hours, then washed and resuspended in calcium free, magnesium free PBS supplemented with FBS. Anti-human CD3 antibody (BioXCell) was added at a concentration of 1 μg/10^6^ cells to avoid potential graft-versus-host disease. Per mouse, 2 × 10^6^ cells in 0.1 mL saline were tail vein injected. Mice engrafted with primary AML cells were sacrificed after 4 weeks. Engraftment was measured by flow cytometry for human CD45+ cells (BD no. 571875).

## Results

To test genetic knockdown of primary human AML and MDS samples, we utilized apheresis products or bone marrow of patients with these malignancies. For analysis of LSC-enriched fractions, specimens were processed as previously described.^6,9,10^ Leukemia stem/progenitor cells (LSPCs) from primary AML patient samples were defined based on their lower reactive oxygen species status as previously published.^6^ LSPCs from primary MDS patient samples were defined by positivity of cell surface marker CD34.^11^ We first sought to optimize electroporation efficiency of LSPCs and ensure electroporation did not affect viability. To do so, we isolated LSPCs from four AML patient samples, followed by electroporation (Figure 1A). For this technique, we adapted the protocol from Brunetti et al.^8^ LSPCs electroporated with a non-targeting siRNA showed no difference in viability compared to a non-transfected control group 24 hours after electroporation (Figure 1B). Next, we tested how effective electroporation was at introducing RNA into the cells. We utilized siGLO Green, a fluorescent oligonucleotide duplex that is commercially available and used as a transfection indicator. As shown in Figure 1C, LSPCs electroporated with siGLO show significant GFP positivity by flow cytometry compared to the non-electroporated cells. This contrasts with the low GFP positivity seen after using the lipofectamine technique (Figure 1D). To further show siGLO is located inside of the cells and translocases to the nucleus after electroporation, we imaged the cells using an ImageStream software during flow cytometry. As shown in Figure 1E, isolated LSPCs from AML patient samples show siGLO in both the cytoplasm and nucleus. Using the ImageStream software, we then quantified how much siGLO translocated to the nucleus in 512 cells. The mean GFP positivity in the nucleus was 60.95% (Figure 1F). Similarly, we used primary MDS patient samples to test the electroporation technique with siGLO. After electroporating a primary MDS bone marrow sample with siGLO as described above, we showed the transfection indicator localizes to the nucleus (Figure 1G). Within the CD34+ stem cell population, the distribution of siGLO transfection indicator in the nucleus is 83.15% (Figure 1H).

**Figure 1.**
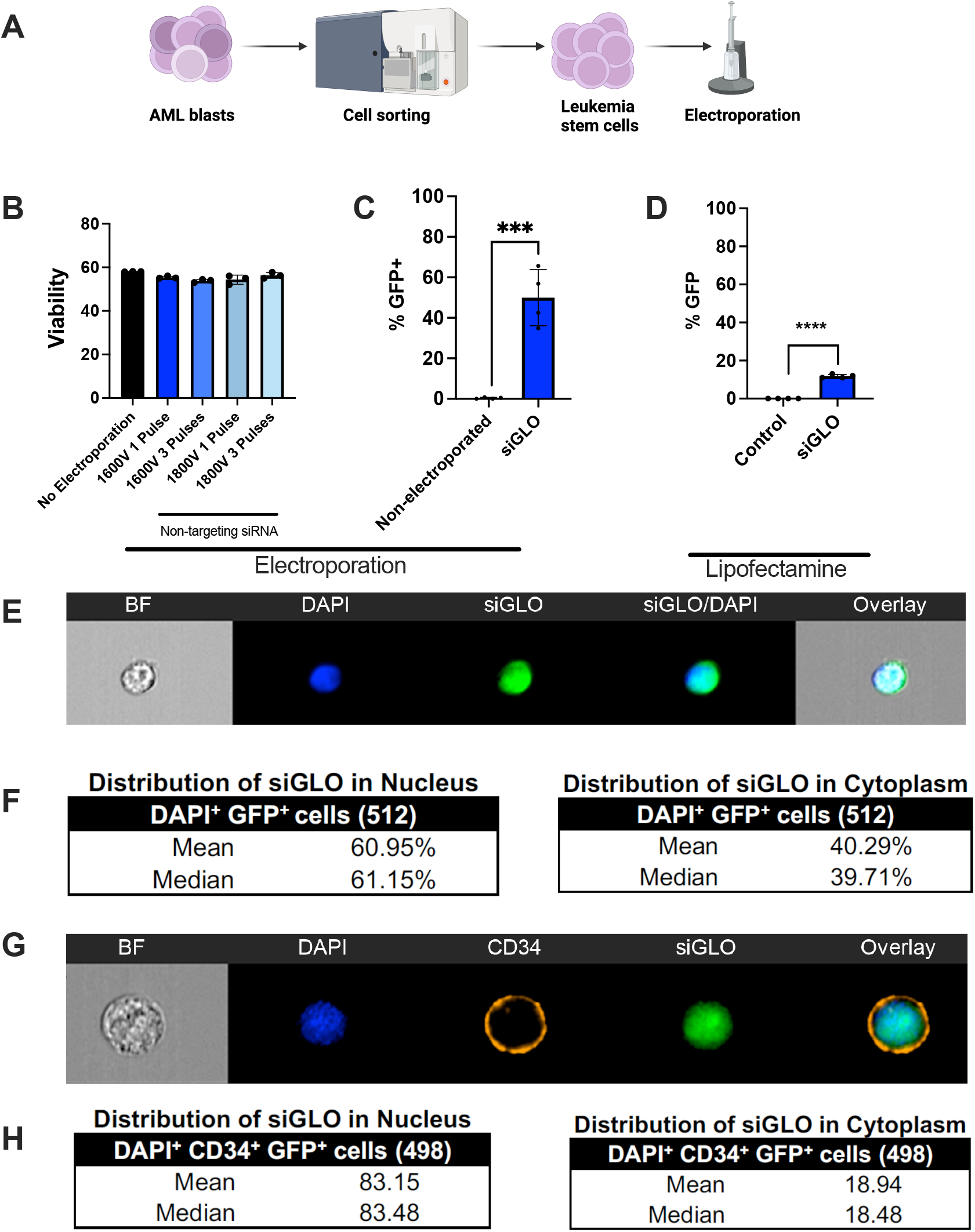
Electroporation leads to effective transfection of leukemia stem cells without affecting viability. (A) Schematic of electroporation technique. Cartoon made with Biorender. (B) Viability of leukemia stem cells 24 hours after electroporation at various voltages as measured by flow cytometry with annexin V and Dapi markers n = 4. (C) Percentage GFP in AML LSCs that were electroporated with siGLO compared to non-electroporated cells as measured by flow cytometry n =4. (D) Percentage GFP in AML LSCs that were transfected with siGLO compared to control via Lipofectamine as measured by flow cytometry. (E) Representative AML LSC electroporated with siGLO and stained with DAPI. (F) Quantification of GFP (siGLO Green) positivity in the nucleus vs cytoplasm of AML LSCs. (G) Representative MDS stem cells electroporated with siGLO and stained with DAPI. (H) Quantification of GFP (siGLO Green) positivity in the nucleus vs cytoplasm of MDS CD34+ stem cells. Statistical analyses were performed using a Student’s t-test. P values are represented as follows: * p ≤ 0.05, ** p ≤ 0.01, *** p ≤ 0.001, **** p ≤ 0.0001. NS = not significant.

To determine whether electroporation of siRNA in LSPCs results in efficient knockdown, we first utilized siRNA against the gene *MYC*. Given *MYC* is critical for leukemia cells, we used this gene as a representative example. We electroporated AML LSPCs with 50nM non-targeting siRNA (siSCR) or 50nM siRNA against the gene *MYC*, respectively. Figure 2A shows decreased MYC expression by qPCR of a representative sample in a time course. As shown in Figure 2B, MYC expression is significantly decreased 48-hours after electroporation of siMYC in LSPCs isolated from 3 AML samples. Next, to examine if our approach would be feasible with a multitude of genes and performed colony forming assays upon gene knockdown, a standard assay to assess leukemia stem/progenitor cell function. We performed siRNA knockdown using the negative control ODC1, which our previous work suggests that loss of this gene alone is insufficient to target primary AML cells^12^ and 7 additional genes within the glutathione S-transferases (GSTs) family known to be expressed in primary human leukemia stem cells (Figure 2B). We suspected that GSTs are critical for LSC function based on previous reports.^10^ Bulk leukemia cells from bone marrow patient samples were electroporated with each target siRNA vs scramble control and were plated in human methylcellulose media. Colonies were counted after 10-14 days in culture, which shows a significant decrease in the number of colonies in cells with various gene knockdowns compared to scramble control (Figure 2C).

**Figure 2.**
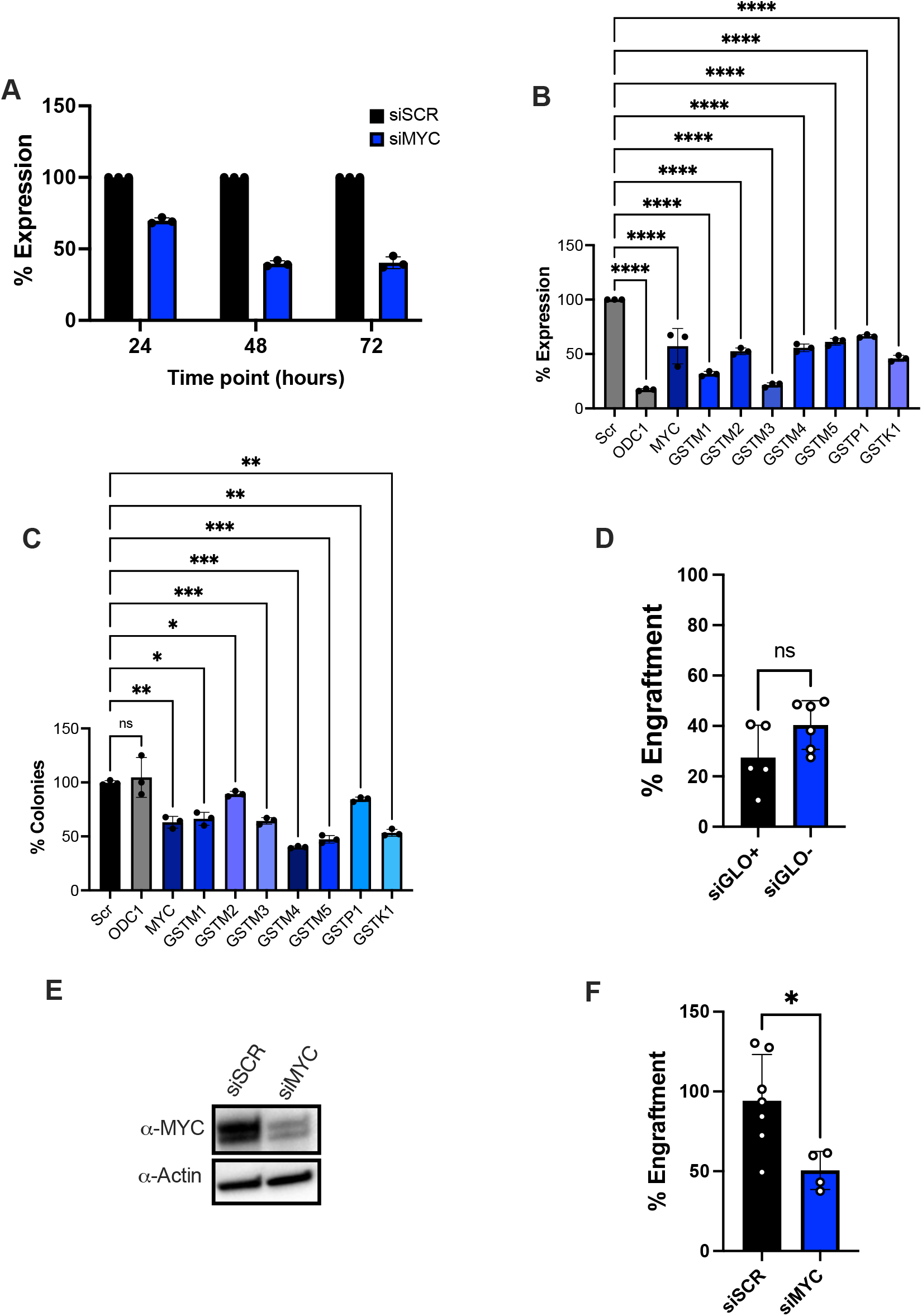
Electroporation leads to effective knockdown and functional readout. (A) RNA expression after MYC knockdown at various time points in LSCs from a representative patient sample measured by qPCR. (B) RNA expression after knockdown of several genes in LSCs isolated from 3 AML patient samples. (C) Percent colonies after knocking down genes from (B). (D) Percent of engraftment of CD45+ human AML cells in NSG-S mice after electroporation of siGLO marker and cell sorting of siGLO positive and siGLO negative cells prior to engraftment in an AML sample. (E) Protein expression of MYC after siRNA knockdown in LSCs from an AML patient sample by western blot. (F) Percent of engraftment of CD45+ human AML cells in NSG-S mice after electroporation with scramble siRNA (siSCR) vs siMYC in an AML sample. Statistical analyses were performed using a Student’s t-test. P values are represented as follows: * p ≤ 0.05, ** p ≤ 0.01, *** p ≤ 0.001, **** p ≤ 0.0001. NS = not significant.

Additionally, we performed patient derived xenograft (PDX) experiments, the gold standard for assessing LSPC activity. To do this, we first tested whether siGLO is effectively introduced into LSPCs. We electroporated LSPCs with siGLO and sorted siGLO positive and siGLO negative cells, followed by injection of each population in conditioned NSG-S mice. As shown in Figure 2D, there was no difference in engraftment, suggesting siGLO does reach LSPCs and these effectively engraft into mice. We next knocked down the gene MYC, a gene known to play a critical role in LSPC function. MYC protein expression tested in a sample of cells was decreased in cells treated with siMYC compared to scramble control (Figure 2E), which led to a significant reduction in tumor burden 4 weeks post transplantation (Figure 2F) indicating effective targeting of the leukemia stem cell population through our methodology. Taken together, our data show we can target effective genetic knockdown in primitive leukemia stem/progenitor cell populations.

## Discussion

In this study, we have demonstrated efficient transfection of genetic material into LSPCs. Genetic manipulation and uptake of genetic material by LSPCs has been historically challenging. Building upon previous CRISPR methods in hematopoietic stem cells (HSCs),^8^ we utilized siRNAs to perform efficient, yet timely genetic knockdown in LSPCs. This is critical as LSPCs have limited viability in culture compared to bulk AML cells or normal HSCs and techniques such as CRISPR which take longer to perturb the genome may not be possible. It is important to note that this technique has certain limitations. First, the electroporation protocol has been optimized for myeloid malignant stem cells, and it is possible other primary cells may require different voltage settings to ensure efficiency without compromising cell viability. Further, knockdown efficiency largely depends on the siRNA. Optimization of the sequence of the gene of interest, siRNA concentration and timeline of knockdown is always recommended for each gene being tested. Additionally, knockdown efficiency depends on both transcript and protein stability of the gene of interest. We have observed here (Figure 2B-E, Supplemental Figure 1) and in previous studies, that the biological effects of knockdown do not necessarily correlate with knockdown efficiency.^9,13-15^ Lastly, although we have shown a very high transfection efficiency in these cells, a minority of cells do not show uptake of genetic material as shown by siGLO. To overcome this, siGLO may be used as a transfection marker to track and sort siGLO+ cells for enrichment of cells with siRNA.

We envision this technique can be used in a several ways to study primary LSPC biology. The effects of a single gene on LSPC activity may be measured through colony forming assays and PDX models post-transfection. Additionally, as shown in Figure 2B, this technique may be used as a rapid, yet efficient screening tool to narrow any targets of interest to further pursue. Furthermore, in our previously published studies, we have shown successful application of this technique for downstream analyses including metabolomics, and pharmacologic studies.^9,13-15^

## Authorship contributions

A.I, H.T, J.R, C.T.J and M.L.A designed research; A.I, H.T, J.R, and M.L.A performed and/or analyzed experiments; A.I and M.L.A prepared the figures; A.I, C.T.J. and M.L.A wrote the manuscript.

## Acknowledgements

A.I. is supported by the National Institutes of Health under Ruth L. Kirschstein National Research Service Award T32CA190216.

C.T.J. is supported by National Institutes of Health (NIH) grants R01 CA200707, R35 CA242376, and P30CA046934; a Leukemia and Lymphoma Society Specialized Center of Research (SCOR) grant; and the Nancy Carroll Allen Endowed Chair.

C.L.J. is support by Leukemia and Lymphoma Society Blood Cancer Discovery Grant, the Rally Foundation, The V Foundation, Alex’s Lemonade Stand Award, and National Institutes of Health (NIH) grant R37CA291896.

M.L.A is supported by BLR&D grant 1IK2BX005603-01A1 from the Department of Veterans Affairs

## References

1. Sekeres MA, Taylor J. Diagnosis and Treatment of Myelodysplastic Syndromes: A Review. JAMA. 2022;328(9):872–880.

2. Shallis RM, Wang R, Davidoff A, Ma X, Zeidan AM. Epidemiology of acute myeloid leukemia: Recent progress and enduring challenges. Blood Rev. 2019;36:70–87.

3. Gentles AJ, Plevritis SK, Majeti R, Alizadeh AA. Association of a leukemic stem cell gene expression signature with clinical outcomes in acute myeloid leukemia. JAMA. 2010;304(24):2706–2715.

4. Eppert K, Takenaka K, Lechman ER, et al. Stem cell gene expression programs influence clinical outcome in human leukemia. Nat Med. 2011;17(9):1086–1093.

5. Hope KJ, Jin L, Dick JE. Acute myeloid leukemia originates from a hierarchy of leukemic stem cell classes that differ in self-renewal capacity. Nat Immunol. 2004;5(7):738–743.

6. Lagadinou ED, Sach A, Callahan K, et al. BCL-2 inhibition targets oxidative phosphorylation and selectively eradicates quiescent human leukemia stem cells. Cell Stem Cell. 2013;12(3):329–341.

7. Lapidot T, Sirard C, Vormoor J, et al. A cell initiating human acute myeloid leukaemia after transplantation into SCID mice. Nature. 1994;367(6464):645–648.

8. Brunetti L, Gundry MC, Kitano A, Nakada D, Goodell MA. Highly Efficient Gene Disruption of Murine and Human Hematopoietic Progenitor Cells by CRISPR/Cas9. J Vis Exp. 2018(134).

9. Pei S, Pollyea DA, Gustafson A, et al. Monocytic Subclones Confer Resistance to Venetoclax-Based Therapy in Patients with Acute Myeloid Leukemia. Cancer Discov. 2020;10(4):536–551.

10. Pollyea DA, Stevens BM, Jones CL, et al. Venetoclax with azacitidine disrupts energy metabolism and targets leukemia stem cells in patients with acute myeloid leukemia. Nat Med. 2018;24(12):1859–1866.

11. Baum CM, Weissman IL, Tsukamoto AS, Buckle AM, Peault B. Isolation of a candidate human hematopoietic stem-cell population. Proc Natl Acad Sci U S A. 1992;89(7):2804–2808.

12. Rondeau V, Berman JM, Ling T, et al. Spermidine metabolism regulates leukemia stem and progenitor cell function through KAT7 expression in patient-derived mouse models. Sci Transl Med. 2024;16(766):eadn1285.

13. Amaya ML, Inguva A, Pei S, et al. The STAT3-MYC axis promotes survival of leukemia stem cells by regulating SLC1A5 and oxidative phosphorylation. Blood. 2022;139(4):584–596.

14. O’Brien C, Ling T, Berman JM, et al. Simultaneous inhibition of Sirtuin 3 and cholesterol homeostasis targets acute myeloid leukemia stem cells by perturbing fatty acid beta-oxidation and inducing lipotoxicity. Haematologica. 2023;108(9):2343–2357.

15. Sheth AI, Althoff MJ, Tolison H, et al. Targeting Acute Myeloid Leukemia Stem Cells through Perturbation of Mitochondrial Calcium. Cancer Discov. 2024;14(10):1922–1939.

