## Supplemental Materials for "siRNA Mediated Genetic Perturbation of Primary Human Leukemia Stem and Progenitor Cells"

### Additional Methods

### Human Specimens

Primary human AML or MDS specimens were obtained by apheresis product, peripheral blood or bone marrow as noted above for corresponding studies. All patients gave informed consent for procurement of samples on the University of Colorado tissue procurement protocol (Colorado Multiple Institutional Review Board Protocol; #12-0173 and #06-0720) or University Health Networks Research Ethics Board #20-5031. The University of Colorado and University Health Network Institutional Review Board approved the retrospective analysis. All specimens were acquired in accordance with recognized ethical guidelines (Declaration of Helsinki and U.S. Common Rule).

### Cell Sorting

Primary human AML specimens were sorted for ROS-low LSCs as previously described.[5] Briefly, specimens were thawed and stained with DAPI (EMD Millipore, no. 278298; dilution 500nM) to exclude dead cells, CD19 (BD, no. 555413; dilution 1:20) and CD3 (BD, no. 557749; dilution 1:40) to exclude lymphocytes, CD45 (BD, no. 571875; dilution 1:40) to identify the blast population and CellROX deep red (Thermo Fisher, no. C10422; dilution 5uM) to identify the 20% of AML blasts with the lowest ROS stain signal, deemed “ROS-Low LSCs”.

### Colony Forming Assays

Primary AML specimens were plated in human methylcellulose (R&D systems) at cell concentrations indicated in respective figures. Samples were plated 2 post-transfection and added into methylcellulose cultures. Colonies were counted at 10-14 weeks after initial plating.

**Engraftment Assays**

All animal studies were done at the University of Colorado under Institutional Animal Care and Use Committee– approved protocol no. 308. The University of Colorado is accredited by the Association for Assessment and Accreditation of Laboratory Animal Care (ALAC), abides by the Public Health Service (PHS) Animal Assurance of Compliance and is licensed by the United States Department of Agriculture.

**Reagents, Antibodies and Primers**

| Reagent | Source | Catalog # or sequence |
|---|---|---|
| MYC siRNA | Dharmacon | L-003282-02-0005 |
| siGLO Green Transfection Indicator | Dharmacon | D-001630-01-05 |
| FITC Mouse Anti-Human CD45 | BD Biosciences | 555482 |
| PE-Cy7 Mouse Anti-Human CD3 | BD Biosciences | 557749 |
| PE Mouse Anti-Human CD19 | BD Biosciences | 555413 |
| PE-CF594 Mouse Anti-Human CD34 | BD Biosciences | 562449 |
| CellROX Deep Red | Thermo Fisher | C10422 |
| PE Annexin V | BD Biosciences | 556421 |
| Beta-Actin Antibody (C4) | Santa Cruz Biotechnology | Sc-47778 |
| Cytokine SCF | PEPROTech | 300-07 |
| Cytokine IL3 | PEPROTech | 200-03 |
| Cytokine FLT3 | PEPROTech | 300-19 |
| Human Methylcellulose Complete Media | R&D Systems | HSC003 |
| Anti-MYC antibody | Cell Signaling | 9402S |

Primer Sequences

| | | |
|---|---|---|
| MYC | AAAGGCCCCCAAGGTAGTTA | GCACAAGAGTTCCGTAGCTG |
| ACTIN | CATGTACGTTGCTATCCAGGC | CTCCTTAATGTCACGCACGAT |
| GSTM1 | GGAGGAACTCCCTGAAAAGC | CGAGAAAATCTACAAAAGTGATCTTG |
| GSTM2 | TGATGTCCTTGAGAGAAACCAA | GGCAGAGATCTTCTCCAAGC |
| GSTM3 | CCAATGGCTGGATGTGAAAT | TCCAGGAGGTAGGGCAGATT |
| GSTM4 | CTTCCGGACCTTGCTCCCTGAA | TGCAACACGGGGACAGCTCA |
| GSTM5 | CACTGCCCCGGTTTTAGTTG | CAGGACTGGGAAAGCATCTG |
| GSTP1 | GGCAACTGAAGCCTTTTGAG | TCAGCGAAGGAGATCTGGTC |
| GSTK1 | TTTGGCTCTGACCGGATG | CACGGCTGGAGGTATAGGG |
| ODC1 | TTTGAGACCAGCCTGGGCAA | TGATCCTCCCACCTCAGCCT |

**Supplemental Figure 1**

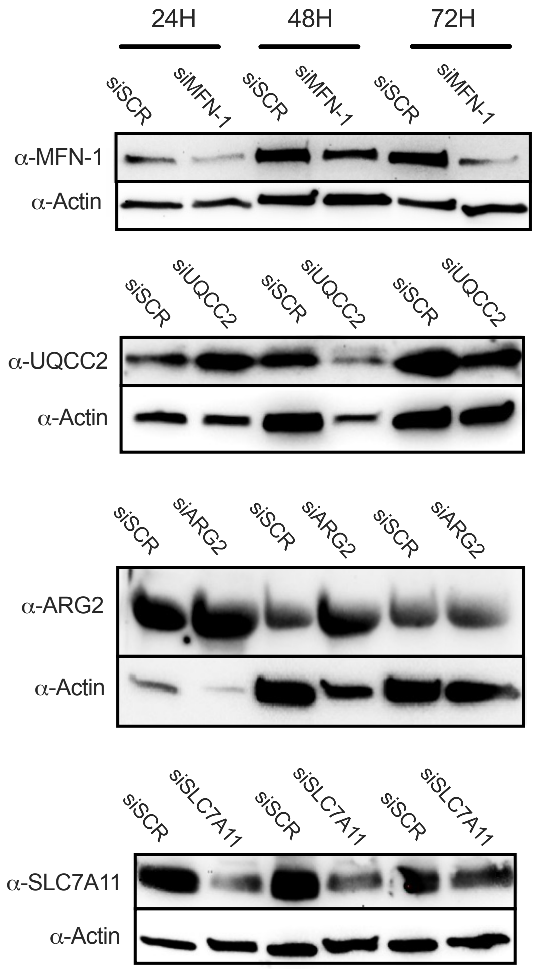


**Supplemental Figure 1**. Protein knockdown occurs at various timepoints. Western blots showing protein knockdown of four different proteins at 24, 48 and 72 hours after LSPCs were electroporated with their respective siRNAs.
